# Electrophysiological Brain Connectivity and Subjective States Evoked by Electrical Stimulation of the Human Mediodorsal Thalamus

**DOI:** 10.1101/2025.11.12.688132

**Authors:** Sofia Pantis, Dian Lyu, Julian Quabs, Weichen Huang, Masaya Togo, Heejung Jung, Eric van Staalduinen, Lindsay Liu Yang, Aidan Chan, Mina Fedor, Robert Fisher, Vivek Buch, Josef Parvizi

## Abstract

Recent advances in human intracranial EEG (iEEG) have enabled new investigations into the role of the thalamus in human brain functions. In this study, we applied direct intracranial electrical stimulation (iES) to the mediodorsal (MD) subregion of the thalamus using both high-frequency (50 Hz, iES_HF_) and low-frequency (0.5 Hz, iES_LF_) procedures to examine its impact on conscious experience and causal brain connectivity in 30 patients with focal refractory epilepsy (128 electrode contacts; 4 ± 1 MD sites per patient). iES_HF_ of the MD elicited reportable changes in conscious experience in 11 of 12 patients (39 sites; 83 stimulations across 27 unique pairs) - predominantly in the visceral, emotional, or somatosensory domains and often described as unpleasant without any lateralization effect. Our connectivity analyses based on iES_LF_ revealed that the cingulate and insular cortices produced stronger electrophysiological responses in the MD (inflow connectivity) than did the sites in the prefrontal cortex (PFC) within the same individuals. Moreover, MD stimulation showed its strongest outflow connectivity to the cingulate, insular, and PFC regions, all significantly stronger than to medial temporal lobe (MTL) structures.

Notably, inflow from both MTL and insula sites to the MD were significantly stronger than their reverse directions, indicating clear asymmetry in connectivity. These findings provide direct evidence that stimulation of the human thalamus can modulate conscious experience. They also highlight the extensive bidirectional connectivity between the MD and cingulate and insular cortices along with asymmetric connectivity between the MD and MTL and insula sites in the human brain.

**SIGNIFICANCE STATEMENT:** Our findings provide a functional and causal map of the mediodorsal thalamus (MD) in the human brain. We provide direct evidence that stimulation of the human thalamus can modulate conscious experience. This study also holds clinical and translational value for identifying thalamic pathways involved in the propagation and generalization of seizures, especially seizures involving the medial temporal lobe, as well as for neuromodulation in epilepsy and other neuropsychiatric disorders, as MD stimulation may not be well-tolerated in human subjects.

## INTRODUCTION

Our understanding of the thalamus is largely based on findings from non-human mammalian brains or group-based neuroimaging studies and causal findings with patients with thalamic lesions, which often extend beyond individual thalamic nuclei or subregions.

The thalamus is known to play indispensable roles in complex cognitive and behavioral functions through reciprocal connections with the cerebral cortex (Parvizi, 2009; Shine et al., 2017). The thalamus is increasingly considered to be highly involved in interregional and network dynamics, serving as a connector hub of various functional networks rather than solely “relaying” information to cortex (Shine et al., 2023). The mediodorsal nucleus of the thalamus in the mid-thalamus (MD) is commonly described as a typical “higher order” thalamic nucleus, mediating cortico-thalamo-cortical communication with the prefrontal cortical (PFC) regions (Sherman, 2007; Saalmann, 2014; Golden et al., 2016; Pergola et al., 2018). The MD has been implicated in many of the PFC functions such as cognitive flexibility, cognitive control, and attention switching (i.e., adapting thoughts and behaviors in dynamic environments) (Golden et al., 2016; Peräkylä et al., 2017; Schmitt et al., 2017; Rikhye et al., 2018; DeNicola et al., 2020); long-term, working, and recognition memory (Watanabe and Funahashi, 2004; Nishio et al., 2014; Danet et al., 2015; Mitchell, 2015; Golden et al., 2016; Peräkylä et al., 2017; Pergola et al., 2018); learning (Mitchell, 2015); decision making (Mair et al., 2015; Mitchell, 2015; DeNicola et al., 2020), and overall information integration and transfer for higher-order cognitive processing (Mitchell, 2015), especially over a temporal delay or when multi-tasking (Pergola et al., 2018). Additional evidence has also found MD involvement in olfaction (Golden et al., 2016) and emotional processing (Golden et al., 2016). Clinically, dysfunction or alterations of the MD have been observed in schizophrenia (Pergola et al., 2015; Pergola et al., 2018; DeNicola et al., 2020), alcohol use disorder (Pergola et al., 2018), Wernicke-Korsakoff syndrome (Mair et al., 2015; Pergola et al., 2018), epilepsy (Patel et al., 1988; Cassidy and Gale, 1998; Bertram, 2014; Zhang and Bertram, 2015; Golden et al., 2016), depression (Cardoso et al., 2009; Li et al., 2010; Li et al., 2024), disorders of consciousness (Maxwell et al., 2004), frontotemporal dementia (Bocchetta et al., 2020), Huntington’s disease (Heinsen et al., 1999), Alzheimer’s disease (Zarei et al., 2010), and Parkinson’s disease (Monje et al., 2020).

Rodent and non-human primate studies utilizing electrophysiology, anterograde and retrograde axonal tracing, and other methods have contributed greatly to mapping thalamic connectivity with the rest of the brain, though the translatability of these findings to human brains has remained to be determined (Mitchell, 2015; Shine et al., 2023). According to these studies, MD projects to PFC, secondary motor area, anterior cingulate, agranular insular, temporal, and parietal cortices and amygdala as well as subcortical areas such as the thalamic reticular nucleus, basal ganglia (including nucleus accumbens) (Steriade et al., 1997; Mitchell, 2015; Golden et al., 2016; Clasca, 2024). Inputs to the MD originate from both cortical (PFC, insular, anterior and mid cingulate, piriform, entorhinal and perirhinal, temporal pole, inferior and superior temporal, subiculum, primary olfactory, and primary and supplementary motor areas) and amygdala, claustrum, midbrain, superior colliculus, ventral pallidum, substantia nigra, hypothalamus, and basal forebrain (Aggleton and Mishkin, 1984; Russchen et al., 1987; Steriade et al., 1987; Yeterian and Pandya, 1988; Gower, 1989; Velayos et al., 1998; Mitchell, 2015; Golden et al., 2016). Noninvasive studies using functional magnetic resonance imaging (fMRI), diffusion-weighted imaging (DWI), and diffusion tensor imaging (DTI) in human participants have revealed similar results, namely, strongest connectivity between MD and the frontal lobe (including PFC, anterior cingulate cortex, motor cortex, and superior, middle, and inferior frontal gyri) as well as temporal lobe regions (including anterior parahippocampal gyrus and amygdala) (Behrens et al., 2003; Zhang et al., 2008; Klein et al., 2010; Jakab et al., 2012; O’Muircheartaigh et al., 2015; Li et al., 2022). Additionally, the two MD subregions are often linked together either via a bridging gray/white matter structure or directly fused together with an inter-thalamic adhesion – both referred to as *massa intermedia* (MI) (Şahin et al., 2023).

Due to the inaccessibility of deep thalamic structures, direct electrophysiological studies of the human thalamus—particularly the mediodorsal nucleus—have been limited (Guye et al., 2006; Evangelista et al., 2015; Filipescu et al., 2019; Chaitanya et al., 2020; Pizzo et al., 2021; Gadot et al., 2022; Soulier et al., 2023; Wu et al., 2023; McGinn et al., 2024), leading to lack of information about the role of this thalamic subregion in conscious human experience and its position within the architecture of whole-brain connectivity. In the present study, we address this gap of knowledge by combining 50Hz electrical stimulation (iES_HF_), to probe a causal link between a given behavioral or subjective state and a given brain region and its anatomical network, and single-pulse electrical stimulation (iES_LF_), also known as cerebro-cerebral evoked potentials (CCEPs), to probe its causal effective connectivity (Parvizi and Kastner, 2018). Our recent publication (Lyu et al., 2025) demonstrated unique feature-specific evoked response patterns across cortical and subcortical regions, and we utilize this method in studying thalamo-cortical connectivity of the MD subregion. Our findings, based on this approach, provide direct evidence that stimulation of the human MD subregion of the thalamus can modulate conscious experience. They also demonstrate the extensive bidirectional connectivity between the MD and the cingulate and insular cortices, alongside asymmetric connectivity patterns with the medial temporal lobe and insula, reflecting the MD’s integrative role within large-scale brain networks.

## MATERIALS AND METHODS

### Participants

The present study includes data from 30 patients (age 20–57, 10 females) selected from a group of more than 250 patients with medically refractory epilepsy who have participated in our intracranial research at Stanford Health Care (SHC) between 2008 and 2025 (Table S1). Patients included in this study had (1) both a high-resolution T1-weighted MRI and a postoperative CT available that could be used for electrode localization, (2) electrical stimulation of at least one kind (iES_HF_ or iES_LF_) conducted, and (3) electrode contacts confirmed to reside within the thalamus by manual review of each patient’s native brain space. Each patient completed an Informed Consent Form approved by the Institutional Review Board at Stanford University prior to participating. All patients willingly chose to partake in electrical stimulation experiments as well as share their data with our research team. No additional electrodes were implanted for solely research purposes. Research in these patients followed the general guidelines detailed in our previous work (Lusk et al., 2025).

### Electrode implantation and localization

A collaborative group at Stanford, including specialists in epilepsy, neurosurgery, psychiatry, neuropsychology, and radiology, determined electrode implantation sites strictly according to each patient’s clinical profile. Only areas potentially involved in a patient’s individual seizure network were implanted. Bilateral depth electrodes through the MD were only implanted if the patient had a clear MI or if the third ventricular transgression between the two thalamic hemispheres was sufficiently small (<3– 4mm). The patients in this cohort were implanted with depth electrodes from Ad-Tech Medical Instruments, with an inter-electrode space of 3–5mm. To localize electrodes, the team used the established iELVis workflow (Groppe et al., 2017). The process began with preprocessing each patient’s T1-weighted MRI using FreeSurfer (https://surfer.nmr.mgh.harvard.edu/fswiki/recon-all), which involved enhancing image contrast, reconstructing the cortical surface in native space, and aligning it to the fsaverage template. Subsequently, the preprocessed MRI was registered to a postoperative CT scan via tools from the FMRIB Software Library (FSL), allowing precise electrode mapping within the patient’s brain space. Electrode contacts were manually annotated in BioImageSuite by identifying bright signals corresponding to electrode locations in the fused MRI/CT images and aligning them with known surgical paths. Electrodes were determined to be localized within the mediodorsal subregion of the thalamus through visual inspection of identifiable anatomical features on the patient’s native space scans.

Additionally, electrode positions were transformed into Montreal Neurological Institute (MNI-305) volume space using an affine transformation, ensuring compatibility with the fsaverage surface and thalamic parcellation. Currently, there are major challenges in identifying specific boundaries of thalamic nuclei in the individual native space, and data from standardized thalamic parcellations were only used as adjunct information. To assess each electrode site’s location within the mediodorsal nucleus, we utilized multiple atlases including Tian(Tian et al., 2020), Brainnetome (Fan et al., 2016), Julich-Brain (Amunts et al., 2020), and “Thalamus Optimized Multi Atlas Segmentation” (THOMAS) (Su et al., 2019). For the former 3 atlases, we used the electrodes’ MNI coordinates and the ICBM 2009c Nonlinear Asymmetric version of the atlases. The THOMAS atlas is an automated pipeline to segment each individual patient’s thalamus according to their own T1 image (https://github.com/thalamicseg/hipsthomasdocker). After the processing, we created a 5 mm sphere around the inner contact centers of mass (i.e., contact neighborhood) and used a binary mask with the spheres to represent the area affected by iES. It then calculated the fraction of voxels in that mask that overlapped with the thalamic nuclei segmentations. We selected 5 mm based on our previous publications, with modifications to account for variations in inter-electrode distances between the prior datasets (Lyu et al., 2025; Togo et al., 2025).

Each atlas’s parcellation of the mid-thalamus differs, with some having distinct “mediodorsal” parcels and others not; as such, each atlas’s count of mediodorsal electrode sites varied. Considering the parcels in each atlas with the highest probability of representing the mediodorsal nucleus, our sample includes 102 mediodorsal sites [anterior division of inferior ventroanterior thalamus (THA-VAia) or posterior division of inferior ventroanterior thalamus (THA-VAip)] according to the Tian atlas, 39 mediodorsal sites [medial prefrontal thalamus (mPFtha)] according to Brainnetome, and 21 mediodorsal sites [mediodorsal nucleus (MD)] according to the Julich Brain atlas. According to THOMAS, 109 electrode sites had a probability between 0.5–100% of being within the mediodorsal nucleus (M = 55.8%, SD = 35.5), with 16 of these sites having a 100% probability. As such, though all MD electrodes included in our sample targeted the mediodorsal nucleus, we can only say with certainty that they were located somewhere within the mid-thalamus.

### Experimental Design and Statistical Analyses

#### Intracranial Electrical Stimulation - High Frequency (iES_HF_)

Direct iES at 50Hz is a standard investigation performed by clinicians during epilepsy monitoring. In this procedure, a pair of adjacent electrodes is stimulated (i.e., *bipolar stimulation*) with square wave electrical pulses. The parameters for iES_HF_ stimulation were 50Hz frequency, 0.2s pulse width, 1–4s duration, and 1 to 6mA current. We followed the methodology outlined in previous publications in regard to experimental procedure, inclusion criteria, subjective response categorization, and quality checking (Duong et al., 2023; Lyu et al.,2023; Duong et al., 2025; Pantis et al., 2025). In brief, we delivered electrical currents to thalamic electrode pairs and asked the patient if they noticed any change in any domain. We noted their verbal responses as well as any relevant facial expressions and movements. As a control, we administered random sham stimulations (i.e., no electrical current) while maintaining identical mannerisms, actions, and probing questions as the genuine stimulations. To ensure the validity and reliability of our findings, we included only responses that occurred without after-discharges, could be replicated with repeated stimulation (unless the patient asked us not to repeat the stimulation due to unpleasant feelings), demonstrated a clear dose-response effect (from lower to higher intensity effects as the current increased), and most importantly, were not present during sham conditions.

#### Intracranial Electrical Stimulation - Low Frequency (iES_LF_)

We administered repeated single-pulse electrical stimulations, a procedure known as cerebro-cerebral evoked potentials (a term we have redefined from the original “cortico-cortical evoked potentials” to encompass the subcortical structures). We used a methodology outlined in a previous publication (Lyu et al., 2025). It is important to note that our analysis of iES_LF_ departs from conventional CCEP method in that it does not depend on an arbitrary threshold for defining significant evoked responses, and instead respects the richness of local field potentials caused in a given region as a result of repeated stimulation of another. Evoked signals often deviate from canonical forms in that they do not necessarily display clear N1 or N2 peaks or their amplitudes do not exceed conventional substantial thresholds. Additionally, previous traditional CCEP studies have typically been conducted in cortical structures and as such the traditional methods may not encapsulate the full complexity of evoked responses involving subcortical regions such as the thalamus.

Electrical pulses were administered at 0.5 Hz in a bipolar configuration (i.e., in pairs of electrodes). Each stimulation pulse lasted 0.2ms and typically involved a current of 6mA, although 4mA was used when needed to reduce seizure risk. Each electrode pair received 42±2 consecutive stimulations. Data that met the following criteria were excluded: (i) recording electrodes within 5mm of the stimulation site, (ii) stimulation or recording electrodes located in white matter, and (iii) two bipolar electrode contacts of either the stimulation or recording pairs located in different anatomical regions.

The iEEG data were acquired at a 1000Hz sampling rate (except for patient 249, for whom data were acquired at 2000Hz sampling rate and down-sampled to 1000 Hz) using the Nihon Kohden system. Preprocessing was performed using a validated in-house pipeline. First, notch filtering was performed at 60, 120, and 180Hz. Next, we excluded noisy channels. Criteria for noisy channels included extreme raw amplitude (>5 SD), frequent spikes (>3× median spike rate), and trial-level metrics such as high mean absolute voltage (>4 SD) or elevated trial variance (>3.5 SD). Trials with mean amplitudes exceeding 4 standard deviations across trials were also discarded. Then, time-frequency analysis was performed using Morlet wavelet decomposition (log-spaced from 1–256Hz, 5 cycles for each wavelet). This generated instantaneous power and phase data downsampled to 200Hz per frequency. Power spectrograms were baseline-corrected, log-transformed, and z-scored across time and frequency. Phase consistency across trials was quantified using inter-trial phase coherence (ITPC), which was square-root transformed and similarly z-scored. Manual inspection was used to validate signal integrity.

We measured the connectivity strength of each stimulation-recording pair using an in-house pipeline based on the power spectrum and ITPC. Specifically, we used a sliding-window cross-correlation to compare the normalized power and ITPC features with a predefined group-derived feature templates from our previous study. Among these features, F1 (Feature 1) is used in the present study. It appears as a sharp wave (increased power in high gamma) with tight phase locking to the stimulation onset (strong ITPC) within the early stage after stimulation (Lyu et al., 2025) (Figure S1). Our previous study showed that F1, defined in the time-frequency domain, is comparable to the time-domain N1 component of the traditional CCEP and may serve as an indicator of feedforward connectivity (Lyu et al., 2025). F1 is also a feature shared between the thalamus-evoked potentials and cortex-evoked potentials which allows us to compare the inflow and outflow connectivity of the thalamus. To measure the F1 connectivity index, Pearson’s correlation coefficient was calculated across time to yield a temporal similarity curve, from which we identified the peak similarity and its timing using MATLAB’s findpeaks function. Connectivity strength was defined as the peak correlation (peak_maxCor) between stimulation–recording pairs (Figure S1).

## RESULTS

We collected data from 128 electrodes (4 ± 1 MD sites/patient) located within the MD (Figure S2). Review of each patient’s T1 MRI revealed presence of MI in 100% of patients. The MI subtypes included adhesion (n = 23), bridge (n = 6), or double bridge (n = 1) as described elsewhere (Şahin et al., 2023).

### Subjective Responses to iES_HF_

The procedure of iES_HF_ was performed in 12 patients yielding data from 39 MD unique sites, stimulated within 27 unique pairs of electrodes, and a total of 83 individual stimulations. Sham stimulations were performed on a random basis 36 times between 10 patients. None of the sham stimulations resulted in subjective responses, yielding a false positive rate of 0% (in line with previous studies supporting the value of sham stimulations as a control) (Fox and Parvizi, 2021). Stimulation of 24 electrodes in 11 of 12 patients evoked subjective responses while stimulation of 3 pairs in only 1 of 12 patients evoked no effect, thus yielding an elicitation rate of 89%.

Site pairs were labeled as eliciting visceral (21 pairs, 9 patients), somatosensory (9 pairs, 5 patients), emotional (4 pairs, 3 patients), and/or auditory/visual (1 pair, 1 patient) effects. As shown in Figure 1 and Table 1, visceral responses were defined as internal bodily sensations involving nausea, dizziness, expansion/contraction, whole-body vibration/shaking, being figuratively awakened/stimulated, lifting, warmth, calm, and euphoria. These effects typically involved the core, throat, head, or whole body.

**Table 1:** Summary of responses to iES_HF_.

| **Patient** | **Laterality of Electrode Pair** | **Response** | **Response Category** | **Response Valence** |
| --- | --- | --- | --- | --- |
| 171 | Bilateral | Fearful/anxious sensation of being stimulated/awakened; heart pumping | Emotion, Visceral | Negative |
| 171 | Right | Stimulated/awakened sensation | Visceral | Negative |
| 178 | Bilateral | Throat contraction, nausea | Visceral | Negative |
| 178 | Right | Nausea | Visceral | Negative |
| 195 | Right | Hallucination of a conversation with a familiar individual | Auditory/Visual | Neutral |
| 197 | Bilateral | Tingling, pressure | Somatosensory | Negative |
| 197 | Bilateral | Tingling | Somatosensory | Negative |
| 211 | Bilateral | Thrill of going down a rollercoaster | Emotion, Visceral | Positive |
| 211 | Bilateral | Soft, warm sensation inside core | Visceral | Positive |
| 212 | Bilateral | Anxiety, fainting/numbing sensation | Emotion, Visceral | Negative |
| 212 | Left | Anxiety, fainting/numbing sensation | Emotion, Visceral | Negative |
| 212 | Right | Fainting sensation, like electricity traveling | Somatosensory, Visceral | Negative |
| 220 | Bilateral | Numbness/tingling, lifting sensation | Somatosensory, Visceral | Negative |
| 234 | Bilateral | Numbness, tingling, expanding sensation, euphoria | Somatosensory, Visceral | Positive |
| 234 | Bilateral | Calm, pushing sensation | Somatosensory, Visceral | Positive |
| 234 | Left | Numbness, calm sensation | Somatosensory, Visceral | Positive |
| 234 | Right | Expanding sensation, euphoria | Somatosensory, Visceral | Positive |
| 237 | Bilateral | Nausea | Visceral | Negative |
| 249 | Left | No effect | No effect | No effect |
| 249 | Bilateral | No effect | No effect | No effect |
| 249 | Right | No effect | No effect | No effect |
| 92b | Left | Numbness/tingling, dizziness | Somatosensory, Visceral | Neutral |
| 92b | Bilateral | Dizziness, nausea | Visceral | Negative |
| 259 | Left | Vibration | Visceral | Neutral |
| 259 | Bilateral | Shaking, nausea | Visceral | Negative |
| 259 | Bilateral | Dizziness, nausea | Visceral | Negative |
| 259 | Right | Dizziness, nausea | Visceral | Negative |

**Figure 1:**
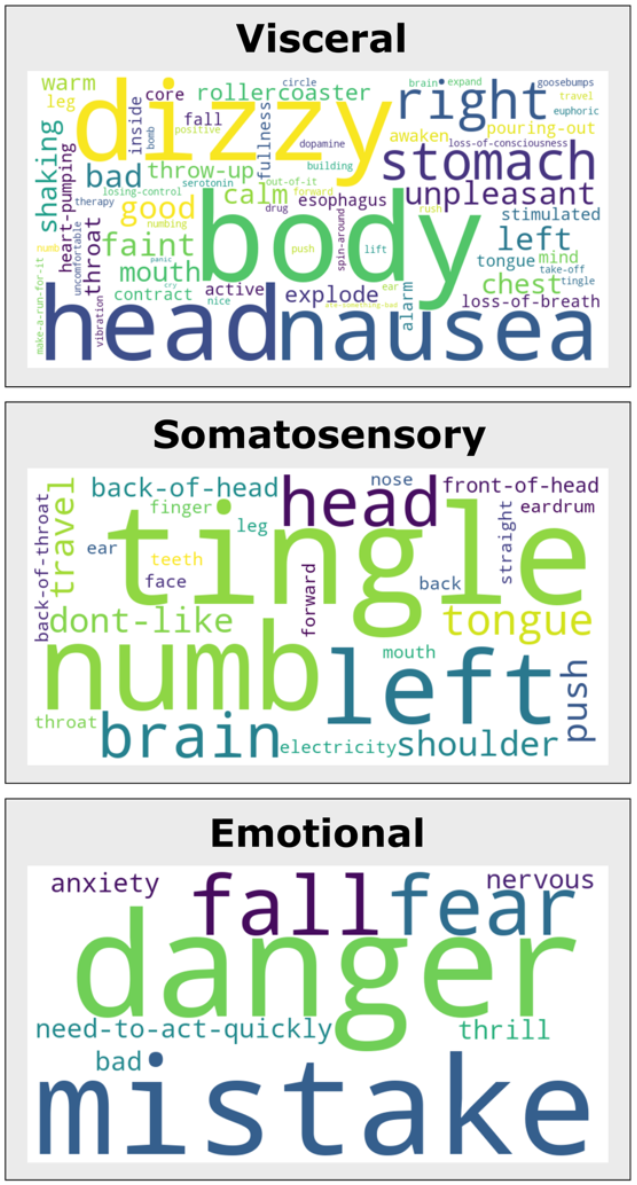
Word cloud representations of verbatim transcriptions of patients’ verbal responses to MD stimulation. Words are separated by category.

Somatosensory responses were defined as tingling, numbness, or pressure (sometimes painful, and sometimes neutral). These somatosensory effects were reported throughout the head, mouth, shoulders, arms, hands, chest, and legs. All somatosensory and emotional responses were reported alongside simultaneous visceral sensation except for two pairs in one patient (Patient 197) who exhibited solely tingling in the head without reporting visceral sensations. Electrical charge delivered in this patient was not higher than those used in the other patients.

Responses to stimulation of a majority of the sites were negatively valenced (15 pairs between 8 patients) in visceral, emotional, or somatosensory domains. These unpleasant subjective changes were reported by the patients as “I felt… like I’m awakened… more active… I think it probably could lead to some danger, some kind of mistake… fear… the need to act quickly… numbness, tingling… almost lifting up out of my head… my brain almost felt like it was going to take off… uncomfortable… really out of it.” By comparison, only few responses were positively valenced. These pleasant changes in patients’ conscious experience were described as “… a feeling in my chest and my stomach, sort of like you are going down a rollercoaster… anxiety that you feel when you are thrilled… soft feeling, warm feeling… as if [a warm blanket] were inside of me… my whole body feels good right now… hard to explain, like you’re on drugs or something… euphoric feeling… yeah, this one feels nice… feels like a serotonin and dopamine just rushed through my whole body and if [the stimulation] was continuous, maybe, it could actually be therapeutic” (of note, the last remark was made by a patient with undergraduate education in neuroscience). Other responses labeled as neutral included mild tingling or vibration. In one single patient, electrical stimulation of MD elicited illusory thoughts of a conversation with a man who was described as Ethiopian (the same as the patient) who was unidentifiable to the patient (“I knew who he is, but I cannot tell who he is right now”). In various stimulations of this electrode pair, the same situation and individual were conjured in her mind, though the conversation topic varied from the Bible to the patient’s epilepsy. The minimum stimulation current that evoked a response was comparable across valence categories, averaging 3.4 mA for negative, 3.0 mA for neutral, and 3.3 mA for positive stimulation pairs. Further, pairs stimulated multiple times at different currents demonstrated clear dose-responsiveness (i.e., greater intensity of response at higher currents).

Our study was not powered to study any lateralization effect. However, we noted that the patterns and valence of responses were similar between left and right hemispheric stimulations. Moreover, no distinct patterns were observed between patients with adhesion/adherent appearance (n = 9) and bridge appearance (n = 3); the one patient with double bridge MI did not undergo the iES_HF_ experiment.

Stimulation of the MI tissue (in patients with bridge forms of it) yielded similar results as the stimulation of the MD site itself. For instance, observations in patient 259 with unusually large ventricles and distinctly large MI provided clear supporting evidence for this claim (see Figure S2 for details).

We are mindful that the precise thalamic nucleus being sampled in every subject could not be definitively determined. However, using probability maps available to us, we analyzed data from a group of subjects in whom the location of electrical stimulation was determined to be within the MD nucleus itself with 100% probability using THOMAS atlas (4 sites in 4 patients). We found no difference in the type of subjective changes reported in these patients compared to the rest of the patients.

### Mapping Causal Connections of the MD with iES_LF_

We collected CCEP data from 28 patients with a combined total of 120 total electrodes within MD. Coverage of the rest of the brain included 3604 unique electrode pairs. We separately analyzed the strength of outflow connectivity (stimulation of MD sites causing evoked responses in other brain sites) and inflow connectivity (stimulation of a given brain site causing evoked responses in MD sites). Our sample totaled 3594 electrode pairs recording CCEPs during MD stimulation (outflow) and 1726 electrode pairs stimulated while CCEPs were measured in MD (inflow) (Figure 2).

**Figure 2:**
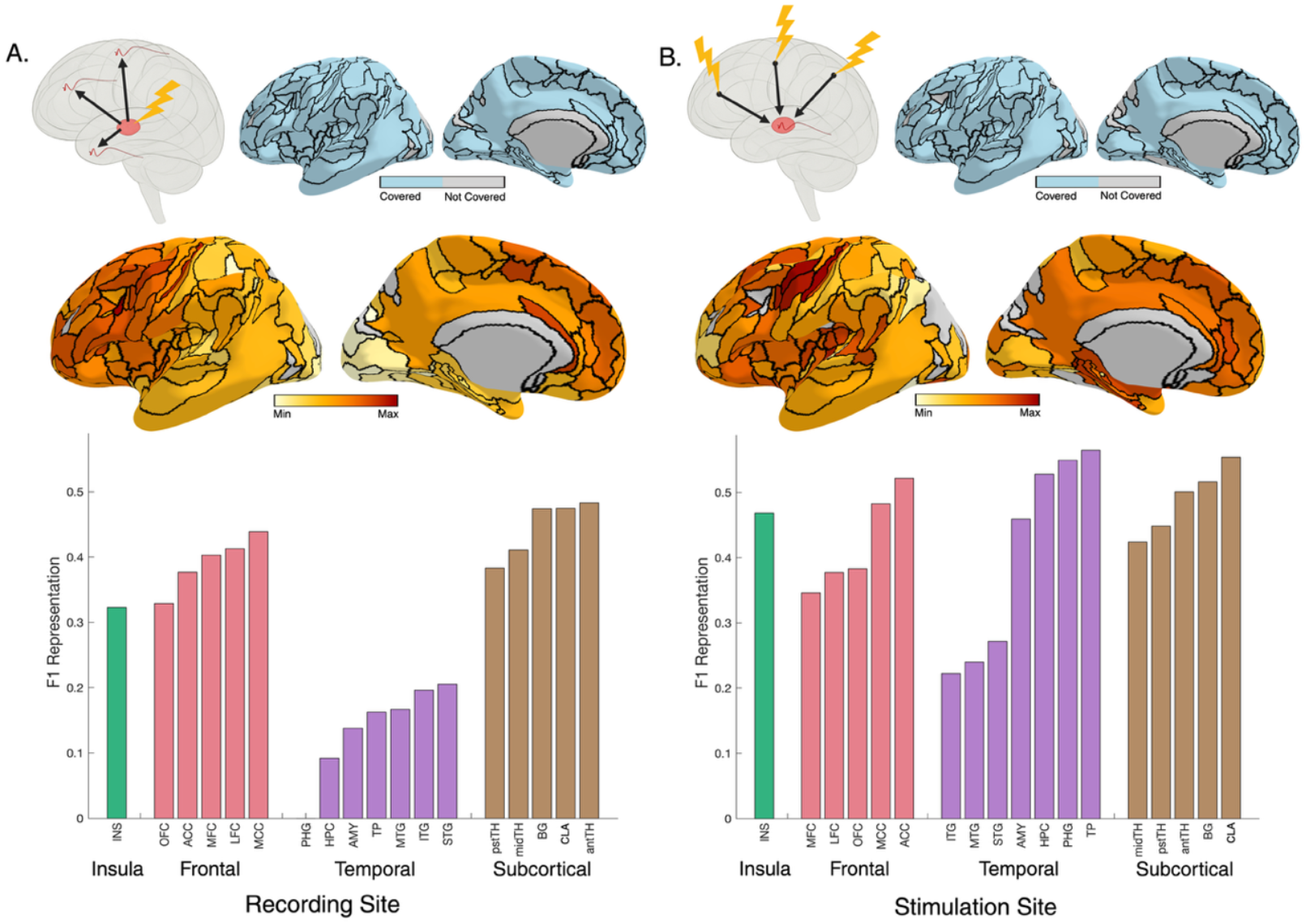
Whole-brain findings from iES_LF_ experiment. Panel (A) presents findings related to outflow connectivity, while panel (B) presents findings related to inflow. The top left of each panel depicts our experimental design, and the top middle and right of each panel depicts whole-brain electrode coverage across all 28 iES_LF_ patients according to Julich-Brain. Below this, we display the mean F1 representation of ipsilateral CCEPs through both anatomical figures (according to both Julich-Brain parcellation) and bar graphs. ACC, anterior cingulate cortex; AMY, amygdala; antTH, anterior nucleus of the thalamus; BG, basal ganglia; CLA, claustrum; HPC, hippocampus; INS, insula; ITG, inferior temporal gyrus; L, left; LFC, lateral frontal cortex; MCC, mid cingulate cortex; MFC, medial frontal cortex; midTH, mediodorsal nucleus of the thalamus (MD); MTG, middle temporal gyrus; OFC, orbital frontal cortex; PHG, parahippocampal gyrus; pstTH, pulvinar (posterior) nucleus of the thalamus; R, right; STG, superior temporal gyrus; TP, temporal pole.

We measured connectivity strength using the F1 feature representation of ipsilateral stimulation-recording electrode pairs (see Methods for details). As shown in Figure 2, regions with strongest outflow connectivity included subcortical structures such as the thalamus (anterior and pulvinar nuclei, as well as other sites within MD), claustrum, and basal ganglia, as well as cortical structures in the insula and frontal lobes including PFC and anterior (ACC) and mid cingulate (MCC) cortices.

We analyzed differences in F1 representation using paired sample t-tests with multiple comparisons corrections at the subject-level for a subset of regions (i.e., regions of interest) where we had the highest electrode coverage, namely, temporal, frontal, and insular cortices. A comparison of outflow connectivity to these ROIs revealed no significant difference between PFC, cingulate, or insula, though each of these regions had significantly greater connectivity strength than both medial (MTL) and lateral temporal lobes (LTL) (Figure 3A-B; see Table S2 for statistics). However, comparing the strength of MD inflow connectivity, using individual patients’ means, suggested cingulate and insula to have significantly greater causal influence over MD (i.e., inflow connectivity) than PFC. Additionally, the inflow connectivity from MTL was significantly greater than the outflow condition. The LTL, however, had significantly less causal influence over MD (i.e., inflow connectivity) compared to all other four regions (Figure 3A-B). Overall, comparing the strength of inflow and outflow connectivity for each cortical region suggested significant asymmetry (i.e., strength of inflow being greater than that of outflow, using subject level data) for only the insula and MTL.

**Figure 3:**
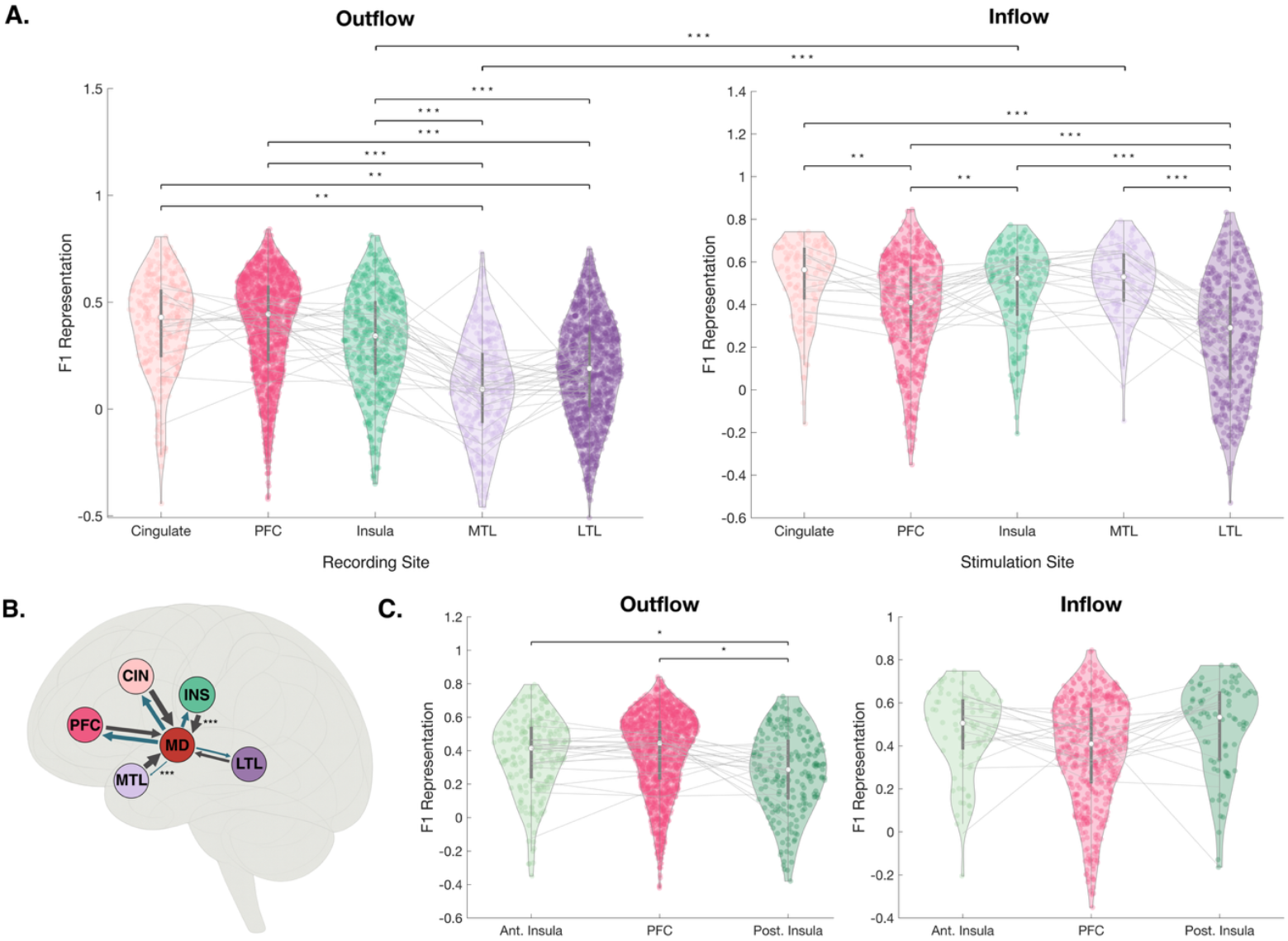
Comparison of MD connectivity between and within cortical ROIs via ipsilateral F1 representation. Panel (A) displays outflow and inflow connectivity patterns between MD and cortical regions of interest, as well as within each region of interest. Panel (B) summarizes Panel (A). Blue arrows indicate outflow and grey arrows indicate inflow. Arrow widths were scaled according to mean F1 values for each condition. Panel (C) compares MD connectivity between PFC, anterior insula, and posterior insula. In (A) and (C), lines between clusters indicate subject-level mean F1 values while lines above clusters indicate a statistically significant difference in F1 values (* indicates p_corrected < 0.05, ** indicates p_corrected < 0.01, *** indicates p_corrected < 0.001 by paired sample t-test with multiple comparisons correction. For detailed statistics results, see Supplementary Table 2 and 3). Cingulate includes anterior cingulate cortex (ACC) and mid cingulate cortex (MCC). Prefrontal cortex (PFC) includes orbitofrontal cortex (OFC), medial frontal cortex (MFC), and lateral frontal cortex (LFC). Medial temporal lobe (MTL) includes amygdala (AMY), hippocampus (HPC), and parahippocampal gyrus (PHG). Lateral temporal lobe (LTL) includes temporal pole (TP), inferior temporal gyrus (ITG), middle temporal gyrus (MTG), and superior temporal gyrus (STG). Anterior insula includes Julich-Brain parcels Id4, Id5, Id6, Id7, Id8, Id9, and Id10. Posterior insula includes Julich-Brain parcels Ig1, Ig2, Ig3, Ia1, Ia2, Ia3, Id1, Id2, and Id3.

Additionally, we examined the differences in connectivity between the PFC and the anterior and posterior subregions of the insula (Figure 3C; see Table S2 for statistics). Posterior insula had significantly lesser outflow connectivity than anterior insula and PFC while all three had equal inflow connectivity.

## DISCUSSION

In the current study, we are reporting two important findings. First, we provide causal evidence that stimulation of the MD subregion of the human thalamus alters conscious experience by eliciting reportable subjective responses. Second, we map the causal effective electrophysiological connectivity (i.e., CCEP) of the MD subregion with cortical and subcortical regions, showing that changes in conscious experience arise not only from the stimulated site itself but from network-level alterations across interconnected brain regions. We found extensive bidirectional connectivity between the MD and both the cingulate and prefrontal cortices, along with asymmetric interactions between the MD and the insula and MTL. Whereas the MD exhibited strong reciprocal causal effective connections with the PFC and cingulate cortex, its connectivity with the insular and MTL structures was markedly asymmetric (Figure 3).

While our findings align with prior work, they reveal some new aspects that might be intriguing.

As noted earlier, according to studies in primates and rodents, MD is connected primarily with the PFC, but also with insular, temporal, and parietal cortices and amygdala. We have previously demonstrated that electrical stimulation of “higher-order” association cortices, such as the PFC, is far less likely to elicit changes in subjective experience, and when such rare changes occur, they tend to be more complex and multimodal in nature (Fox et al., 2020; Pantis et al., 2025). In this context, the finding that stimulation of the MD elicited changes in approximately 90% of instances—predominantly within visceral domains—is intriguing and likely indicates that this thalamic subregion has direct access to cortical and/or subcortical regions involved in the integration of visceral signals. Conceptually, our findings add intriguing causal evidence linking a *subcortical* structure such as the thalamus with human conscious experience (Parvizi, 2009; Parvizi et al., 2022). In keeping with this, lesion studies dating back to the 1950s demonstrated that bilateral mediodorsal thalamotomy could relieve psychiatric symptoms or unfavorable affective symptoms like anxiety, obsessive-compulsive behavior, aggression, depression, and emotional reactiveness to schizophrenic hallucinations (E. A. Spiegel, 1951, 1953; Spiegel and Wycis, 1968). A recent lesion study also posits the MD’s causal role in psychosis (Pines et al., 2025).

Our connectivity results also provide novel information about the asymmetry, and possibly directionality, of information processing through MD. Greater inflow, compared to outflow connectivity, from MTL and the insula was present along with reciprocally symmetric and strong connections between MD and PFC and cingulate, while the LTL had little connectivity with the MD without any obvious asymmetric directionality. Two previous publications in rodent (Shyu et al., 2004) and rabbit (Sikes and DeFrance, 1985) brains also reported activation of the ACC upon stimulation of the MD. Our observation of greater outflow connectivity from MD to anterior insula than posterior insula aligns with previous reports of iES in anterior insula eliciting primarily visceral and emotional responses (Duong et al., 2023). Additionally, though previous studies have found interconnection between MD and the primary olfactory cortex, lateral occipital cortex, primary auditory cortex, and motor cortex, these connections were relatively weak according to our analysis, and thus it is unsurprising that we did not witness any olfactory or motor responses (and only one auditory/visual response) in the iES_HF_ experiment (Russchen et al., 1987). The present work is the first study of human MD connectivity using electrophysiological measures, and as such, our data offer valuable insights that other non-invasive human connectivity studies lack due to methodological limitations. The information obtained from our observations provides new insights into the placement of the MD subregion of the human thalamus within the architecture of brain connectivity.

Lastly, in the clinical context, it is important to be aware of the high prevalence of unfavorable visceral and emotional responses elicited by the stimulation of the human MD subregion since thalamic DBS is now commonly considered as a target for therapeutic interventions in patients with epilepsy or neuropsychiatric disorders (Elias et al., 2020; Ilyas et al., 2022). This is particularly important because MD is adjacent to the anterior nucleus of the thalamus (ANT), and a superior approach to ANT may locate the deepest contacts in MD. Of note, a prior study using DBS-style MD stimulations (145Hz) demonstrated significant disruptions in working memory task performance, though subjective states of patients were not probed (Peräkylä et al., 2017). It remains to be determined if cognitive impairment caused by MD stimulation could have been biased by unpleasant and possibly distracting feelings induced by electrical stimulation. Moreover, understanding the connectivity of the MD (and its connection with the contralateral thalamus through the MI), particularly their relationships with MTL as highlighted in our study, has important implications for identifying thalamic pathways involved in the propagation and generalization of seizures originating from the MTL. For instance, the MI may serve as a critical bridge through which seizures in one hemisphere can spread to the contralateral side via white matter conduction and/or gray matter ephaptic spread between thalami; furthermore, though studies traditionally refer to MI as a grey matter structure, probabilistic DTI and tractography reveal white-matter tracts passing through MI that connect bilateral limbic, frontal, and temporal regions, supporting its potential as a commissural conduit within the limbic network (Kochanski et al., 2018; Borghei et al., 2021).

In closing, while our study revealed important and unique electrophysiological and causal information about the human MD, we are mindful that our approach suffered from several limitations. First, our electrode coverage was limited by clinical necessity. As a result, we could not probe the connectivity of the human MD with regions that are rarely implanted for epilepsy monitoring, such as the brainstem or occipital and parietal cortices. We were also unable to attribute our findings to specific thalamic nuclei for several reasons. For instance, the electrical fields generated by our stimulations could have surpassed the boundaries of mediodorsal nucleus, and currently there is lack of information about the size of the electrical fields generated with single pulse stimulations of macro-electrodes used in epilepsy practice.

Moreover, there are significant variations among subjects and, more importantly, these variations are not homogeneous in the three dimensions of the thalamus (Morel et al., 1997). Our present results in clinical subjects lack the required microstructural specificity to support or reject claims from extant literature about varying connectivity profiles and functional maps between different cytoarchitectonic subdivisions of the MD. In the future, as imaging technologies continue to advance, we envision a time when thalamic electrode locations can be mapped with far greater precision allowing the connectivity of the MD and the subjective correlates of stimulating specific cytoarchitectonic regions of the thalamus to be revealed in unprecedented detail. Such fine-grained insights are now beginning to emerge for cortical regions (Duong et al., 2023), and the same resolution may soon extend to the thalamus.

## Acknowledgements

The authors are grateful to the patients who took part in this research as well as the medical staff at Stanford Health Care, from nurses to patient safety monitors to EEG technicians to epilepsy fellows and attendings, that made this research possible. This work was supported by a research grant from the US National Institute of Neurological Disorders and Stroke (R01NS137650) to J.P.

## Notes

**Conflict of interest statement:** The authors declare no competing interests.

### Competing Interest Statement

The authors have declared no competing interest.

