## Supplementary Materials for "Electrophysiological Brain Connectivity and Subjective States Evoked by Electrical Stimulation of the Human Mediodorsal Thalamus"

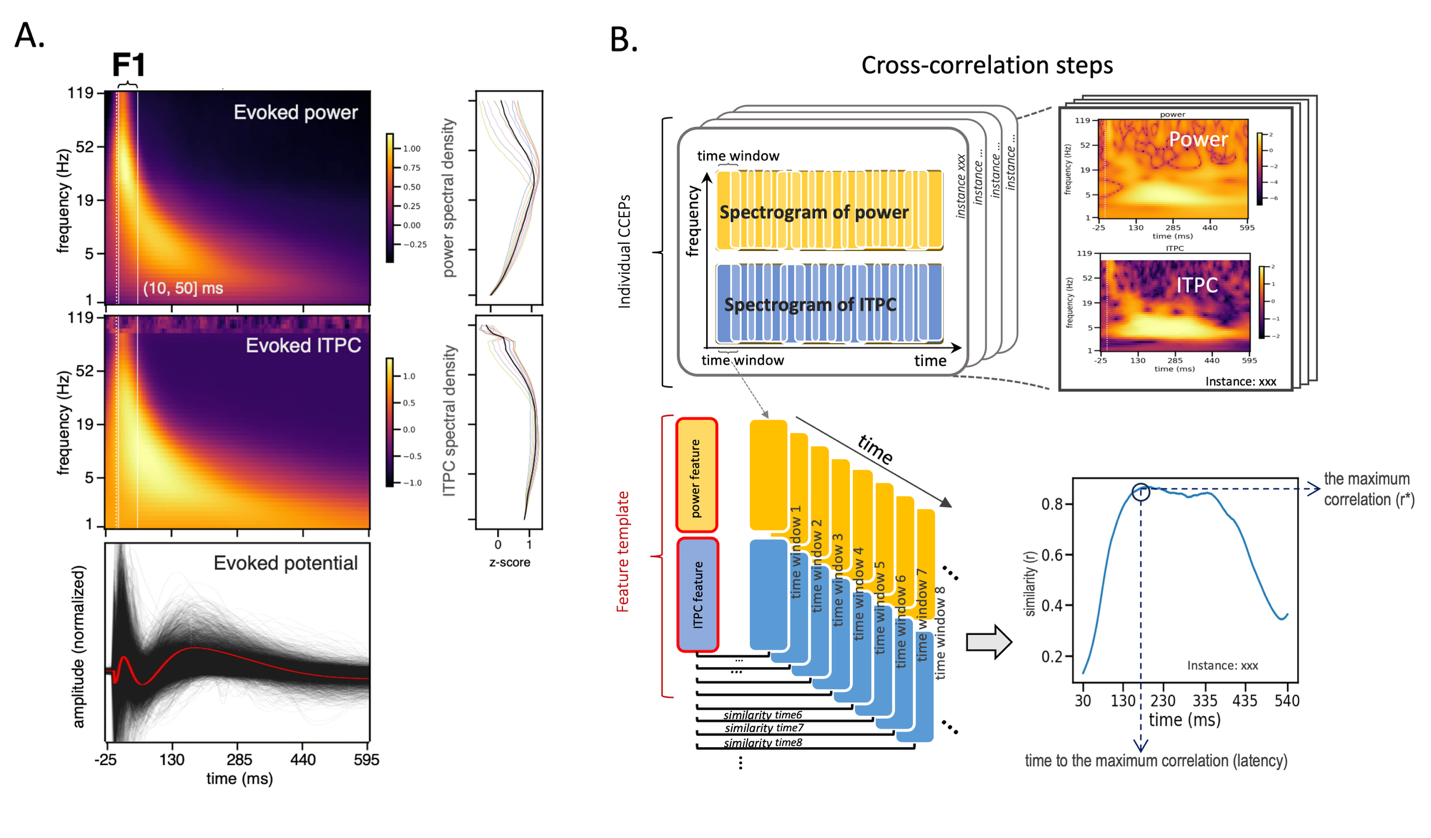

**Figure S1: Electrophysiological neural features and decoding process for Feature 1 (F1).** (A) Feature 1 shown in the power (top), ITPC (middle) and time (bottom) domain. The feature was discovered using the evoked power and ITPC of whole-brain stimulation and recording, which was found to be shared between the thalamus and cortex evoked potentials. For visualization purposes, the group-averaged ipsilateral cortico-cortical evoked potential data was presented as F1 is shown to be the strongest there. The power and ITPC spectral density in the early stage after stimulation was shown at the right (colored lines indicate individual time points, and the black line indicate the average among them), with the y-axis shared with the spectrograms to the left. (B) Sliding-window cross correlation steps to calculate the feature representation in individual CCEPs.

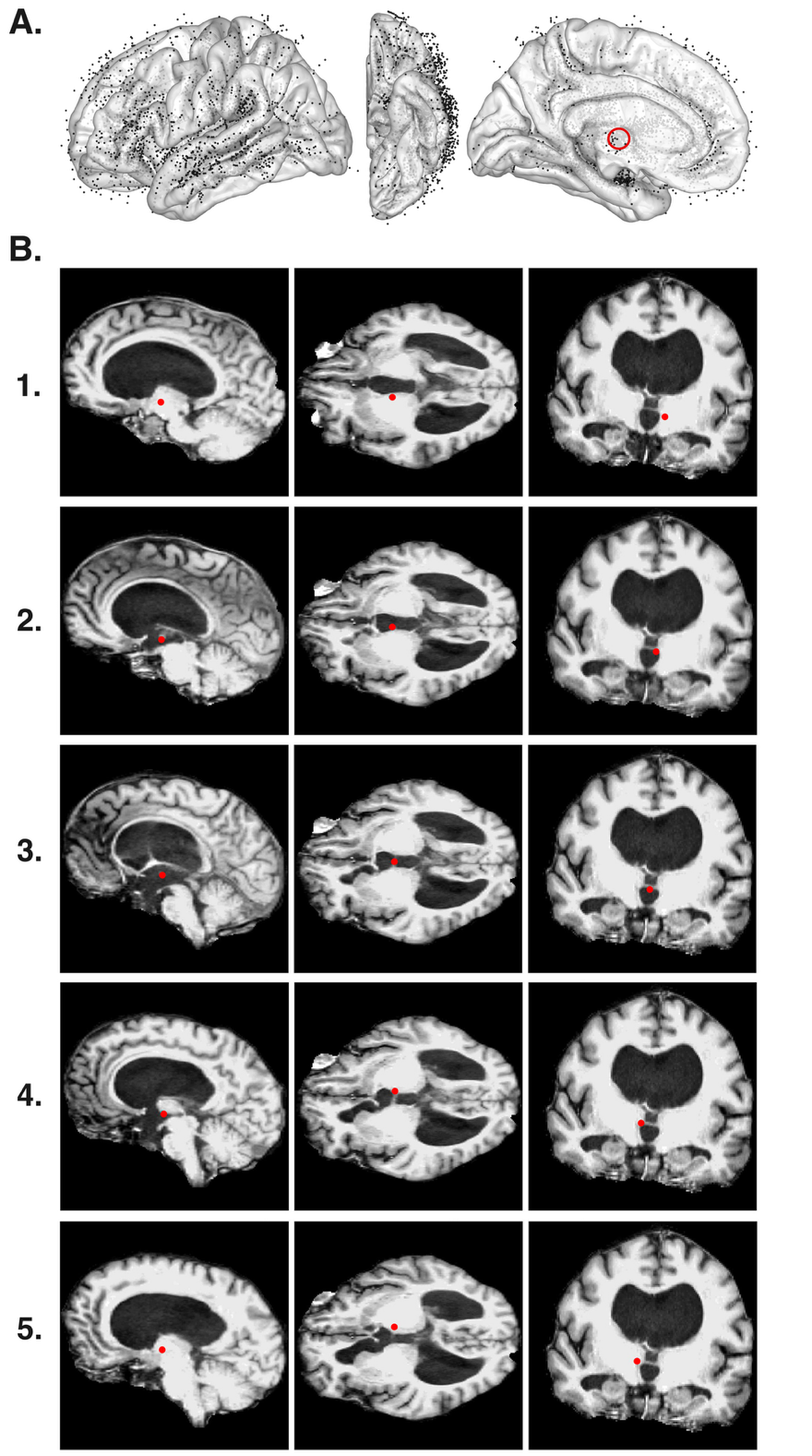

**Figure S2: Electrode coverage.** (A) All electrode sites included in our study are shown in lateral, inferior, and medial views of an MNI brain - totaling 128 MD electrodes (4 ± 1 MD sites per patient) and 4695 non-MD electrodes (M = 168 ± 39 sites per patient). The red circle denotes the approximate location of the MD. Electrodes appearing outside the brain were located within the brain in the native anatomical space. *When electrode locations are transferred from native to standard anatomical space, shifting in their anatomical location may occur. All electrodes are projected onto the left hemisphere.* (B) Example MD electrode sites from patient 259 are shown in this patient’s native space. This patient had unusually large ventricles, allowing for clear observation of her massa intermedia (MI), which appears as an inter-thalamic bridge structure.

**Table S1**: **Patient demographics and experiment participation.** A = Asian; Ambi = Ambidextrous; B = Black; F = Female; H = Hispanic; L = Left; M = Male; NHPI = Native Hawaiian or Pacific Islander; R = Right; W = White

| **Patient** | **Sex** | **Age** | **Race/Ethnicity** | **Handedness** | **Number of MD Electrode Contacts** | **iES_LF_** | **iES_HF_** |
| --- | --- | --- | --- | --- | --- | --- | --- |
| 169 | M | 23 | W | R | 8 | Yes | No |
| 170 | M | 40 | A | R | 4 | Yes | No |
| 171 | M | 52 | A | R | 3 | Yes | Yes |
| 172 | M | 47 | A | R | 5 | Yes | No |
| 178 | M | 28 | W | R | 4 | Yes | Yes |
| 181 | F | 23 | W | R | 5 | Yes | No |
| 182 | F | 37 | W | R | 4 | Yes | No |
| 190 | F | 50 | W | R | 2 | Yes | No |
| 195 | F | 47 | B | R | 5 | Yes | Yes |
| 196 | F | 40 | W | R | 5 | Yes | No |
| 197 | M | 20 | B | R | 4 | Yes | Yes |
| 198 | M | 38 | B | R | 6 | Yes | No |
| 199 | M | 36 | W | R | 4 | Yes | No |
| 203 | M | 51 | H | R | 4 | Yes | No |
| 208 | M | 57 | H | R | 5 | Yes | No |
| 211 | F | 32 | NHPI | R | 5 | Yes | Yes |
| 212 | M | 35 | H | R | 5 | Yes | No |
| 213 | M | 20 | W | R | 4 | Yes | No |
| 214 | F | 28 | H | R | 4 | Yes | No |
| 219 | F | 27 | B | R | 4 | Yes | No |
| 220 | M | 32 | W | R | 4 | Yes | No |
| 229 | M | 31 | W | Ambi | 5 | Yes | Yes |
| 231 | M | 22 | H | R | 2 | Yes | No |
| 234 | M | 27 | W | L | 4 | Yes | No |
| 237 | M | 29 | W | R | 2 | Yes | Yes |
| 239 | F | 50 | W | R | 3 | Yes | Yes |
| 245 | M | 45 | W | R | 3 | Yes | Yes |
| 249 | M | 40 | W | R | 7 | Yes | Yes |
| 92b | M | 45 | H | R | 3 | No | Yes |
| 259 | F | 35 | H | R | 5 | No | Yes |

**Table S2: Statistics for Figure 3.** Includes t and p-values [Benjamini-Hochberg (BH) corrected for multiple comparisons] from paired sample t-tests with multiple comparisons corrections conducted on the subject-level average F1 values in each region.

| **Condition** | **Comparison** | **t** | **p_corrected** |
| --- | --- | --- | --- |
| Outflow | Cingulate - PFC | -0.23 | 0.822 |
|  | Cingulate - Insula | 1.08 | 0.332 |
|  | Cingulate - MTL | 4.25 | 0.001 |
|  | Cingulate - LTL | 4.26 | 0.001 |
|  | PFC - Insula | 2.08 | 0.068 |
|  | PFC - MTL | 6.52 | < 0.001 |
|  | PFC - LTL | 8.19 | < 0.001 |
|  | Insula - MTL | 6.68 | < 0.001 |
|  | Insula - LTL | 4.20 | 0.001 |
|  | MTL - LTL | -1.98 | 0.073 |
|  | Ant. Insula - PFC | -0.93 | 0.364 |
|  | PFC - Post. Insula | 3.03 | 0.027 |
|  | Ant. Insula - Post. Insula | 2.59 | 0.023 |
| Inflow | Cingulate - PFC | 3.51 | 0.006 |
|  | Cingulate - Insula | -0.11 | 0.912 |
|  | Cingulate - MTL | -0.12 | 0.912 |
|  | Cingulate - LTL | 6.25 | < 0.001 |
|  | PFC - Insula | -3.23 | 0.006 |
|  | PFC - MTL | -2.12 | 0.069 |
|  | PFC - LTL | 4.70 | < 0.001 |
|  | Insula - MTL | 0.74 | 0.586 |
|  | Insula - LTL | 9.36 | < 0.001 |
|  | MTL - LTL | 5.04 | < 0.001 |
|  | Ant. Insula - PFC | 1.26 | 0.337 |
|  | PFC - Post. Insula | -1.42 | 0.337 |
|  | Ant. Insula - Post. Insula | 0.22 | 0.832 |
| Cingulate | Outflow - Inflow | 1.42 | 0.178 |
| PFC | Outflow - Inflow | 1.45 | 0.178 |
| Insula | Outflow - Inflow | 4.72 | < 0.001 |
| MTL | Outflow - Inflow | 6.37 | < 0.001 |
| LTL | Outflow - Inflow | 1.63 | 0.178 |
